# Role of the obligate STRIPAK complex component Mob4 in zebrafish vascular development and stability

**DOI:** 10.64898/2025.12.16.694419

**Authors:** Tvisha Misra, Shimon M. Rosenthal, Mengyi Song, Nathan J. Stutt, Laura McDonald, Ashish R. Deshwar, Anne-Claude Gingras, Ian C. Scott

## Abstract

The **STR**iatin **I**nteracting **P**hosphatases and **K**inases (STRIPAK) complex is a conserved multiprotein module with functions in cell proliferation, cell growth, and cancer. Primarily examined through proteomic and *in vitro* studies, there is a relative lack of insight into the *in vivo* function of the STRIPAK complex during development. Motivated by the prominent interaction of STRIPAK and the CCM3 adapter protein, which is associated with the vascular disease cerebral cavernous malformation (CCM), we sought to examine the role of the conserved STRIPAK component Mob4 in early zebrafish vascular development. We use proximity-dependent biotinylation (BioID) coupled with mass spectrometry to show that Mob4 is a component of the STRIPAK complex in zebrafish. Loss of *mob4* leads to pronounced cardiovascular and neuronal defects, including a leaky cerebral vasculature, hypersprouting of venous endothelial cells and poor neuronal branching. Conditional transgenic models further demonstrate a continuous and broad requirement for Mob4 in development. Taken together, these results suggest a key role for STRIPAK function in vascular development and stability, replicating hallmarks of CCM.

## Introduction

The coordination of signalling pathways is a key process during development and physiological homeostasis whose deregulation has been linked to a variety of conditions, including vascular diseases. Multiple signalling events are mediated through scaffolded protein complexes, which are thought to provide specificity to substrate selection and spatio-temporal control of the signalling events. The <u>Str</u>iatin-interacting <u>p</u>hosphatase <u>a</u>nd <u>k</u>inase (STRIPAK) complexes are large, highly conserved multiprotein structures scaffolded by the large Striatin proteins (Castets et al., 1996; Goudreault et al., 2009). Striatin orthologs and multiple STRIPAK components are conserved through evolution (three in mouse and humans, and one each in *0. melanogaster* and *C. elegans* (Hwang and Pallas, 2014). In mammals, paralogous expansion and mutually exclusive interactions among STRIPAK components give rise to multiple distinct assemblies, although the specific functions of these complexes remain incompletely understood. As its name suggests, the STRIPAK complex contains both phosphatases (the PP2A phosphatase catalytic and scaffolding subunit, the latter of which directly binds to Striatin) as well as kinases (the GCK-III family of Sterile-20 kinases that are recruited to the STRIPAK scaffold via the adapter protein CCM3/PDCD10) (Kean et al., 2011; Moreno et al., 2000). STRIPAK also associates with other kinases to modulate various signalling pathways, including the Hippo pathway (Couzens et al., 2013; Ribeiro et al., 2010). Consistent with these roles, dysregulation of Striatin family members and their interactors has been implicated in human disease, particularly in cancer cell migration and metastasis (Bisoyi et al., 2022; Hwang and Pallas, 2014; Kim et al., 2020; Kuck et al., 2016; Madsen et al., 2015; Qiu et al., 2020; Shi et al., 2016; Tang et al., 2020).

Striatins are important for various processes such as dendrite outgrowth, hearing development, dorsal closure, excretory canal formation, sarcomere formation, and neuronal development (Bartoli et al., 1998; Berger et al., 2022; Lahav-Ariel et al., 2019; Lant et al., 2015; Li et al., 2018; Nadar-Ponniah et al., 2020). Given the cell-type-specific expression and potentially redundant or specialized roles of Striatin paralogs, further work is needed to assess the developmental roles of the STRIPAK complex. Among the many STRIPAK components is MOB4 (formerly known as Mob3 or Phocein;(Baillat et al., 2001)), a member of the evolutionarily conserved monopolar spindle-one-binder family best known for its role as a kinase regulator. The presence of multiple paralogs for most STRIPAK components in vertebrates complicates comprehensive functional analyses of the complex. However, MOB4 is consistently identified as a core STRIPAK component in multiple vertebrate species (Baillat et al., 2001; Goudreault et al., 2009; Jeong et al., 2021) and human MOB4 is able to functionally replace its *0rosophila* counterpart (Schulte et al., 2010). Rat Mob4 is enriched in brain tissue in dendritic and Purkinje cells and to a lesser extent in several other tissues (Baillat et al., 2001; Bailly and Castets, 2007; Castets et al., 1996; Haeberle et al., 2006). In *0rosophila* S2 cells, Mob4 localizes to mitotic centres and kinetochores and regulates spindle organization in mitotic cells (Trammell et al., 2008). Loss of Mob4 leads to defects in spermatid differentiation (Santos et al., 2023), axonal transport, and microtubule organisation (Schulte et al., 2010; Sepp et al., 2008). In *C. elegans*, *mob-4* is widely expressed, with loss leading to reduced lifespan and thermotolerance (Jahan et al., 2021). In planarians, Mob4 knockdown loss leads to increased body size and defects in proportionality and tissue rescaling during regeneration (Schad and Petersen, 2020). In zebrafish, Mob4 is localized to Z discs in the sarcomere and is required for sarcomere organization, with further roles in neuronal and cartilage development, cell survival and organization of microtubule networks (Berger et al., 2022). Despite these insights, a comprehensive analysis of MOB4 and STRIPAK function during early vertebrate development remains lacking.

Here, we describe the role of Mob4 in early zebrafish development. Using proximity-dependent biotinylation, we define the *in vivo* interactome of Mob4, which closely parallels previously reported *in vitro* data from mammalian systems, confirming its role as a core STRIPAK component. Analysis of a *mob4* mutant model reveals defects in cardiac, vascular and neuronal development. In the vasculature, Mob4 loss leads to defects in blood vessel integrity and endothelial cell migration, specifically in venous endothelial cells. We show that *mob4* mutant defects can be rescued by ubiquitous expression of a Mob4 transgene and find that Mob4 is required continuously for proper vascular development. Via restoration of Mob4 function, we show that *mob4* phenotypes can be partially rescued by re-expression of Mob4. Taken together, these results indicate broad and continuous requirements for STRIPAK/Mob4 in vertebrate development and suggest a critical role for the STRIPAK complex in vascular development and stability.

## Results

### Mob4 is a primary component of the STRIPAK complex in zebrafish

To define the *in vivo* interactome Mob4, we used a biotin-dependent proximity labelling approach employing the biotin ligase TurboID (Branon et al., 2018) in zebrafish embryos (Rosenthal et al., 2021) (Fig 1A). We injected one-cell stage embryos with mRNA encoding Mob4, N-terminally tagged with TurboID and C-terminally tagged with EGFP (TurboMob4EGFP). TurboEGFP RNA and uninjected embryos were used as controls. Embryos were incubated in biotin-supplemented egg water from 12 to 48 hours post-fertilization (hpf), followed by protein purification and mass spectrometry analysis (Rosenthal et al., 2021). Data from 2 replicates of *TurboMob4EGFP* were compared against multiple replicates of *TurboEGFP* and uninjected controls using the SAINTexpress software (Teo et al., 2014). Proteins with a Bayesian false discovery rate (BFDR) ≤5% are considered high-confidence proximity partners (a complete list of proximal interactors and their human orthologs is provided in Supplementary Tables S1 and S2). As expected, the top enriched hits largely comprised components of the STRIPAK complex, including Striatin/Striatin3/Striatin4, Slmap, Fam40a/Strip1, and Cttnbp2/Cttnbp2L (Fig 1B). Gene Ontology (GO) gene set enrichment analysis of the human orthologs of high-confidence hits showed the “STRIPAK complex” as the most significantly enriched GO term (Fig 1C, and Supplementary Tables S3). The *in vivo* Mob4 interactome therefore strongly suggests a conserved role for Mob4 in STRIPAK complex function during zebrafish embryogenesis.

**Figure 1:**
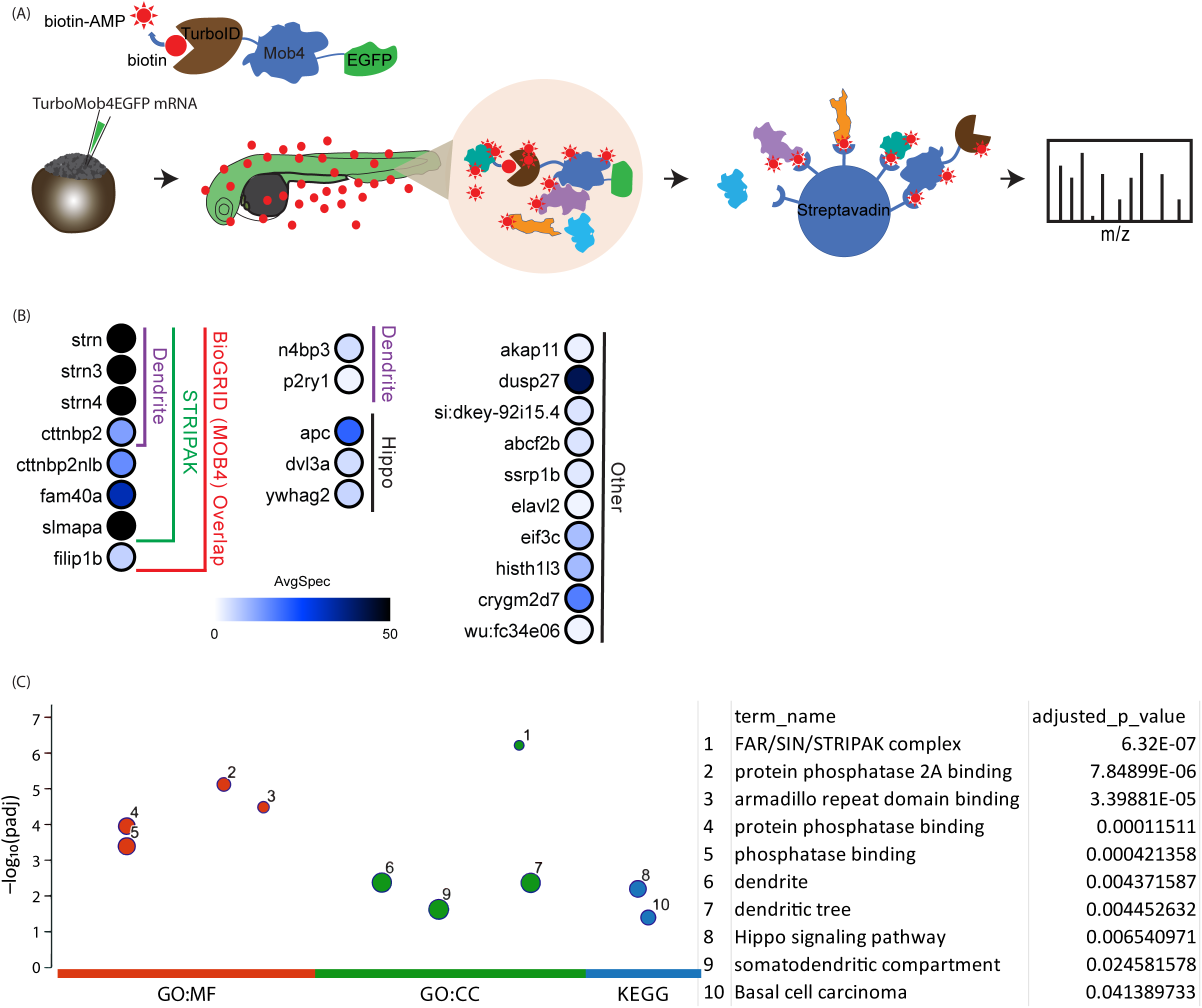
*In vivo* interactome of Mob4 in zebrafish embryos. (A) Schematic of pipeline to conduct BioID *in vivo*. *In vitro* transcribed *TurboMob4EGFP* mRNA is injected at the single cell stage. Embryos are transferred to 800µM biotin containing egg water at 16 hpf and collected for protein purification at 48 hpf. Biotinylated proteins are enriched on streptavidin beads and further analysed with mass spectrometry. (B) Proteins with a BFDR ≤5% after SAINT analysis are shown. The average abundance (by spectral count) is visualized by the colour of the dot on the dot plot. (C) Gene Ontology (GO) term enrichment analysis results for the Human orthologues of the high confidence (BFDR ≤5%) hits.

### *mob4* **loss leads to early cardiovascular defects**

To investigate the role of Mob4 in development, we generated a *mob4* mutant model via injection of CRISPR/Cas9 ribonucleoprotein (RNP) complexes targeting exon 3 (Fig 2A, red circle). An allele with an 11 base pair deletion was isolated (Fig 2B, top panel *mob4 mutant*), leading to a premature STOP codon early in the kinase regulation domain (Fig 2A). *mob4* mutant embryos exhibited cardiac oedemas at 56 hpf (Fig 2C’, black arrowhead), indicating early cardiovascular defects. We use double transgenic *kdrl:EGFP*; *gata1:dsRed* embryos to visualize endothelial cells (ECs) with EGFP and red blood cells with dsRed. In contrast to wild-type embryos, in *mob4* mutant embryos, blood flow was lost over time, as evidenced by stationary blood cells in the tail trunk (Fig 2D and D’, white arrowhead). Additionally, *mob4* mutant embryos exhibited instances of cranial haemorrhaging (Fig 2D’, yellow arrowhead) as seen by aggregation of dsRed-positive blood cells in the hindbrain. The severity of both cardiac oedemas and cranial haemorrhages in *mob4* mutant embryos at 72 hpf varied (Fig 2E-E’’’, magenta and blue asterisks, respectively). Haemorrhages were not observed in the tail trunk vasculature, suggesting a vascular bed-restricted role for Mob4.

**Figure 2:**
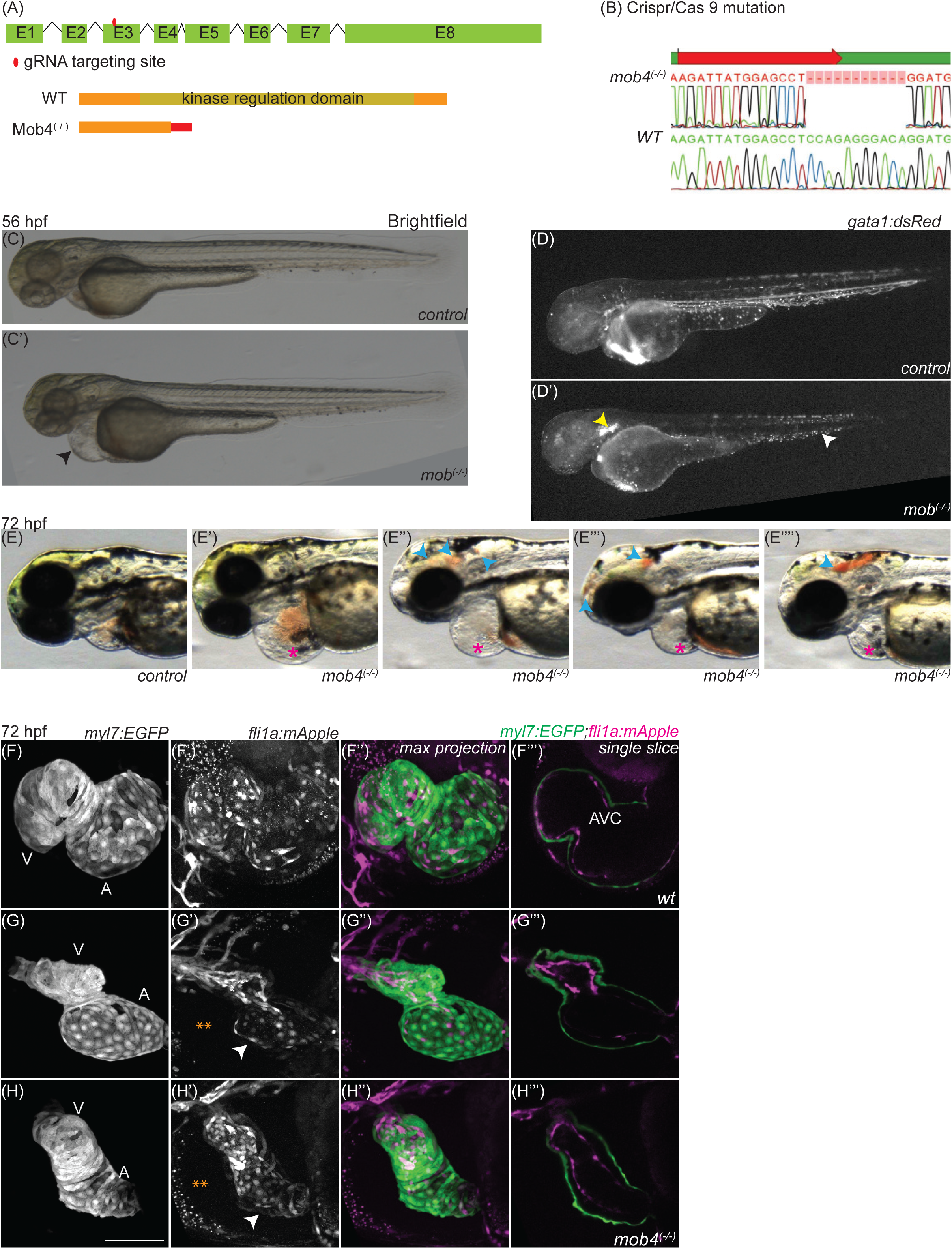
Crispr/Cas9 mediated Mob4 lof leads to cardiovascular defects. (A) Schematic of *mob4* locus with gRNA targeting site (top panel) and corresponding WT protein and mutant peptide expected (bottom panel). Red rectangle in *mob4* mutant peptide represents non conserved residues induced by the mutation before eventual STOP codon. (B) Sequence traces of 11bp deletion in *mob4* mutant allele (top panel) compared to *WT* sequence (bottom panel). (C-C’) Brightfield images of *control* (C) and *mob4 mutant* (C’, black arrowhead: cardiac oedema) embryos at 56 hpf. (D-D’’) *gata1:dsred* labelled blood cells in *control* (D) and *mob4* mutant (D’, yellow arrowhead: haemorrhage, white arrowhead: loss of blood flow). (E-E’’’’) Representative brightfield images of degrees of cranial haemorrhaging (blue arrowheads) and cardiac oedemas (magenta asterisks) at 72 hpf (E’-cardiac oedema no visible haemorrhage, E’’-cardiac oedema + cranial haemorrhage E’’-mild cardiac oedema + two fore- and hindbrain haemorrhages, E’’’-slight cardiac oedema + large hindbrain haemorrhage). (F-H’’’) heart morphology in 72 hpf *control* (F-F’’) and *mob4 mutant* (G-G’’, H-H’’) embryos. A: Atrium. V: Ventricle. Orange asterisks: oedema pericardial space. White arrowheads: AVC, Atrioventricular canal. I’’’, J’’’, K’’’: single slices of z-stacks showing endocardium (*kdrl:hRasmCherry*) and myocardium (*myl7:EGFP*). Scale bar = 100um.

Given the cardiac oedemas and progressive loss of blood flow in *mob4* mutant embryos, we next examined cardiac development via the use of *kdrl:mApple; myl7:EGFP* transgenic embryos, in which the endothelium and endocardium are labelled with mApple and the myocardium is labelled with EGFP (Fig 2F-H’’’). While atrial and ventricular chambers and a patent outflow tract with an endocardial lining were apparent in *mob4* mutants at 72 hpf, in many cases, heart looping and morphogenesis of the atrioventricular canal were abnormal (Fig 2H-H’’’, orange asterisks). Large oedemas were apparent in the majority of *mob4* mutant embryos (Fig 2H-H’’’, white arrowheads), which may contribute to defects in cardiac development. Overall, overt defects in cardiac and vascular development were apparent in *mob4* mutants.

### *mob4* mutants exhibit defects in cerebral vasculature

Given the cerebral haemorrhaging observed in *mob4* mutants, we next examined the development of the cranial vasculature. At 48 hpf, the overall architecture of the vasculature was not affected in *mob4* mutants, with major vessels and structures such as the dorsal midline junction (DMJ) easily identifiable (Fig 3A-B’’). Haemorrhages could be readily identified by the accumulation of blood cells outside the vasculature at 48 hpf (Fig 3B-B’’, yellow arrowhead. By 96 hpf, vascular defects were more pronounced in *mob4* mutants (Fig 3C-D’’). Blood flow was largely halted, as evidenced by stationary blood cells. Further, collapsed vessels were evident such as the Dorsal Longitudinal Vein and Dorsal Midline Junction (Fig 3D’’, asterisks). We next used time-lapse imaging to track the development of cerebral haemorrhages and cessation of blood flow in *mob4* mutants starting at 54 hpf in the *kdrl:EGFP*; *gata1:dsRed* double transgenic background. Fig 3E-F shows projections of z-stacks of single time points from imaging over 54 to 72 hpf (Supplementary movie 1). Angiogenesis could be observed in the extension of ECs to establish new central arteries (Fig 3E, asterisks), which subsequently lumenized and established blood flow. Early cerebral blood flow was established in *mob4* mutant embryos (Fig 3F-F’’’). However, in some cases (arrowheads Fig 3F’’’-F’’’’’) after a nascent vessel was lumenized and blood flow established, a haemorrhaging event occurred with blood leakage. Blood vessels continued to experience blood flow after haemorrhage events. Similar time-lapse imaging was carried out on the tail trunk vasculature, beginning at 54 hpf when vessels were lumenized and robust blood flow could be observed in not only the larger dorsal artery (DA) and caudal venous plexus (CVP) but also the smaller intersegmental vessels (ISVs) (Supplementary movie 2). The extent of tail trunk blood flow in *mob4* mutant embryos at 54 hpf varied broadly, with either robust flow at 54 hpf or a relative absence of blood cells and flow from as early as 54 hpf (Fig 3G-I). In *the mob4* mutant embryos, blood flow was established in early development, similarly to control embryos, but was gradually lost over time, though the exact developmental stage at which this happened varied between embryos. Independent of the degree of blood flow, *mob4* mutants exhibited aberrant sprouting of trunk blood vessels, which arose from either ISVs (red asterisks) or the ventral vascular bed (blue asterisks) (Fig 3G-I), with aberrant vascular sprouting occurring in a manner seemingly independent of loss of flow in affected vessels (Fig 3I). Defects were evident in retinal vasculature as well (Fig 3J-J’’).

**Figure 3:**
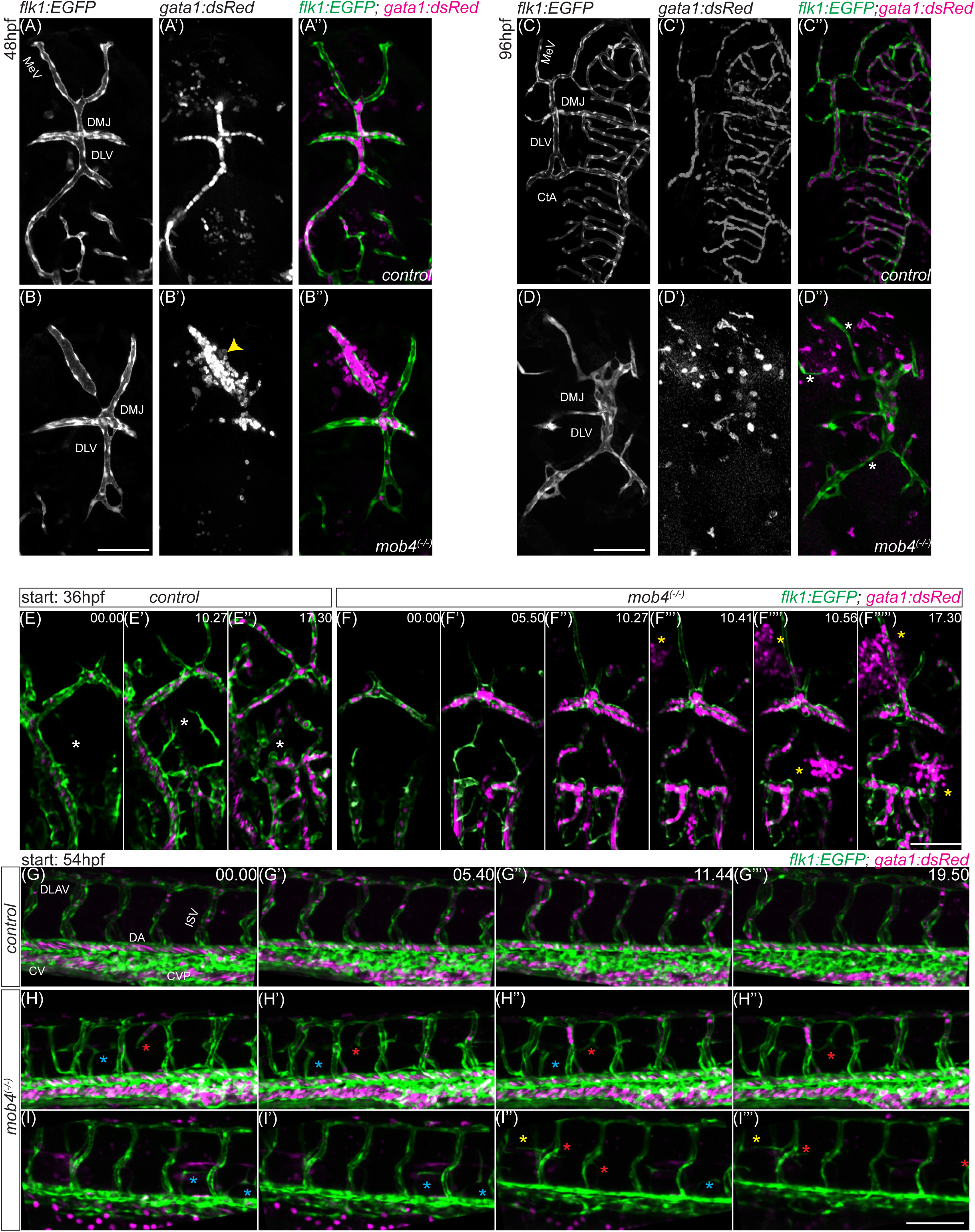
Mob4 lof leads to cranial haemorrhages and vascular defects: (A-D’’) Cranial vasculature (*kdrl:EGFP*) and blood cells (*gata1:dsRed*) in *control* (A-A’’: 48 hpf, C-C’’: 96 hpf) and *mob4 mutant* embryos (B-B’’: 48hpf, D-D’’ 96hpf); Yellow arrowhead: haemorrhage, white asterisks: collapsed blood vessels). (E-F’’) Snapshots of time lapse videos of cranial vascular development and blood flow from 36 hpf in *WT*(E-E’’) and *mob4 mutant* (F-F’’). Yellow asterisks: Haemorrhages. (G-I’’) Snapshots of time lapse videos of tail trunk vasculature development and blood flow from 36 hpf in *WT*(G-G’’) and *mob4 mutant* (H-H’’, I-I’’) embryos. Over sprouting endothelial cells are indicated with asterisks across time; each colour follows a single vessel. DMJ: dorsal midline junction, MsV: Mesencephalic Vein, ISV: intersegmental vessels, DA: dorsal aorta, CV: caudal vein, CVP: caudal venous plexus.

The blood vessel comprises the inner endothelial layer composed of ECs and surrounding supporting architecture consisting of basal, lamina pericytes and astrocytes (Gaengel et al., 2009). Defects in pericyte localization and function lead to vascular instability and haemorrhaging (Rajan et al., 2020). We found that pericyte localization (as indicated by use of a *pdgfrb:Citrine* transgenic line (Ando et al., 2016; Vanhollebeke et al., 2015) was not obviously affected in *mob4* mutant embryos compared to *control* embryos (Fig S1); with both head and tail trunk vasculature comparably decorated with pericytes. Overall, while the development of the cerebral vasculature was initiated normally in *mob4* mutants, there was a subsequent failure in vascular integrity prior to a loss of blood flow. While leakage was not observed in the tail trunk vasculature, abnormal angiogenesis did similarly result from the loss of Mob4.

### *mob4* mutants exhibit hypersprouting of venous endothelial cells in the tail trunk

We next examined the nature of tail trunk EC hypersprouting in *mob4* mutant embryos. Primary angiogenic sprouting of ECs from the dorsal aorta (DA) along the myotome boundaries occurs from 20-36 hpf. Subsequently, during secondary angiogenesis, venous ECs from the posterior cardinal vein (PCV) migrate along the same myotome boundaries from 36-60 hpf. The intersegmental vessels thus formed are only functionally determined as arterial or venous depending on the interaction of the newly formed secondary vessels with the previously established primary network (Isogai et al., 2003). As our initial analysis of *mob4* mutants suggested defects in secondary angiogenesis (Fig 3H-I’’’), we used the Notch reporter line *tp1:EGFP* to label arterial ECs (Quillien et al., 2014). In double transgenic *tp1:GFP*; *kdrl:hRasmCherry* embryos all ECs are labelled with membrane-targeted mCherry and arterial ECs are labelled with GFP (Fig 4 A-C’’). In *mob4* mutant embryos, hypersprouting ECs (Fig 4 B’, C’, yellow asterisks) were labelled with mCherry but negative for EGFP, suggesting a non-arterial nature. Time-lapse imaging of embryos from 56-71 hpf was carried out to visualise EC hypersprouting in real time. Representative wild type and *mob4* mutant embryos are shown in Supplementary movie 3. As expected, hypersprouting was already evident in some *mob4* mutant embryos at 56 hpf (Supplementary movie 3C-C’’ white asterisks); however, subsequent *de novo* hypersprouting events were also observed (Supplementary movie 3B-B’’, yellow asterisks). Hypersprouting ECs were all negative for EGFP. We further used a *mrc1a:GFP* transgenic line to mark venous and lymphatic ECs, (Fig 4 D-D’’) (Jung et al., 2017). In *mob4(-I-)*; *kdrl:hRasmCherry*; *mrc1a:EGFP* embryos, all hypersprouting ECs in the tail trunk were labelled with both EGFP and mCherry. Thus, our data strongly suggest that hypersprouting ECs do not arise during the primary angiogenic wave but are instead venous in origin.

**Figure 4:**
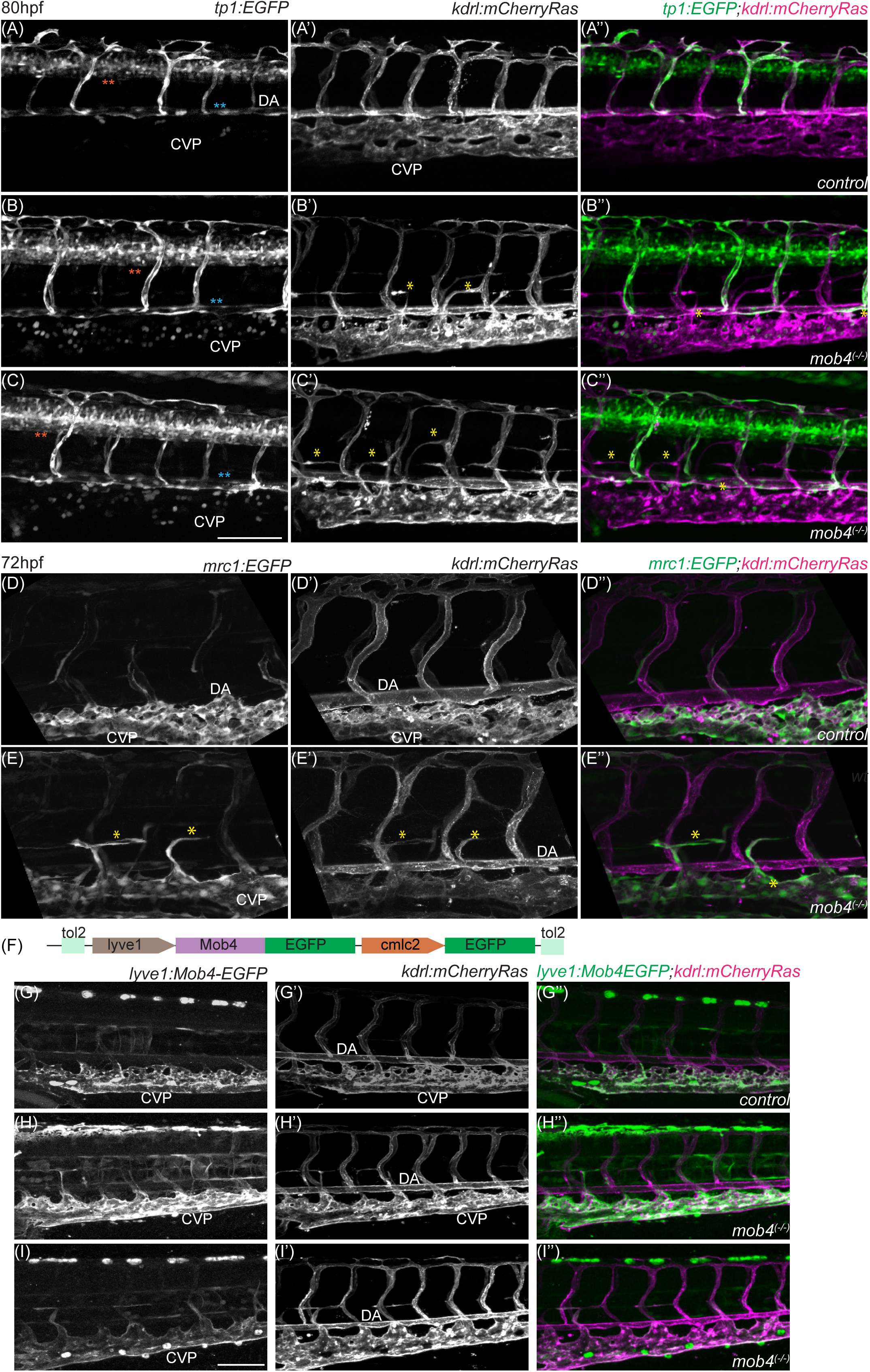
Mob4 lof leads to defects in sprouting of lymphatic derived vessels. (A-C’’) Tail trunk vasculature (*kdrl:hRasmCherry*) and notch signalling reporter (*tp1:GFP*) in *control* (A-A’’) and *mob4 mutant* (B-B’’, C-C’’) in 80 hpf embryos. mCherry positive and GFP negative hypersprouting ECs are marked with yellow asterisks. Cyan asterisk: DA, orange asterisk: notochord. (D-E’’) Tail trunk vasculature in double transgenic (*kdrl:hRasmCherry;mrc1:EGFP*) to label all ECs with mCherry and lymphatic ECs are cells in *control* (D-D’’) and *mob4 mutant* (E-E’’, C-C’’) in 72 hpf embryos. Cherry and GFP double positive hypersprouting ECs are marked with asterisks. (F) Schematic for transgenic to express Mob4EGFP in lymphatic cells. (G’-J’’’)_*wt* (G-G’’) *mob4 mutant;kdrl:hRasmCherry;(lyve1:Mob4EGFP;cmlc2:EGFP)* and Mob4 loss (I-J’’) *mob4 mutant;kdrl:hRasmCherry;* (*lyve1:Mob4EGFP;cmlc2:EGFP)* embryos at 3.5dpf. Scale bar-100µm. CVP: Caudal Venous Plexus, DA: Dorsal Aorta

We next asked if the hypersprouting EC phenotype can be rescued by the expression of Mob4 in venous ECs. *Lyve1* is a highly specific marker of lymphatic cells, although it is also active in a subpopulation of venous endothelial cells. Zebrafish *lyve1* expression is observed in both lymphatic and venous vascular structures such as the caudal vein, common cardinal vein, and the posterior cardinal vein (Okuda et al., 2012) (Fig 4G-G’’). Strikingly in *mob4(-I-)*; *kdrl:hRasmCherry*; *lyve1:Mob4EGFP* embryos, there were no discernible instances of hypersprouting blood vessels and vessels of the tail trunk were comparable to those of control embryos (Fig 4G-I’’). Thus, we conclude that *lyve1* promoter-driven *mob4* re-expression results in rescue of hypersprouting of venous/lymphatic ECs from the caudal venous plexus. This indicated a cell-autonomous role for Mob4 in venous/lymphatic ECs during the secondary angiogenic phase of vascular development.

### *mob4* mutants exhibit neuronal branching defects

Given the role of many STRIPAK complex members in neuronal development and function, we next examined the developing nervous system in *mob4* mutant embryos. Using a pan-neuronal *Tg[ngnl:mKate]* (Kinney et al., 2020) line to image neurons, we first examined *mob4* mutants at 84 hpf. Both impaired arrangement of cranial neuronal bundles and reduced sprouting of tail trunk motor neurons were consistently observed (Fig 5A-B’’). To further examine the nature of neuronal defects, time-lapse imaging was employed from 54 hpf, when motor neuron migration from the spinal cord is initiated (Myers, 1985; Myers et al., 1986). In *mob4 mutant* embryos, initial motor neuron migration/outgrowth at 54 hpf was impaired (Fig 5D-D’’, blue and yellow asterisk) compared to wild type (Fig 5C-C’’). Primary outgrowth of motor neurons is followed by secondary and tertiary outgrowth (Fig 5C’’-C’’’, red, green and purple asterisks follow single motor neurons). In contrast, in *mob4* mutants, various instances of migration defects were observed: impaired (yellow asterisk) or no (blue asterisk) secondary migration, looping of axonal outgrowths (brown asterisk) or short non-elongating outgrowths (yellow asterisk) (Fig 5D-D’’). These results show that complete Mob4 loss leads to severe defects in the development of the nervous system.

**Figure 5:**
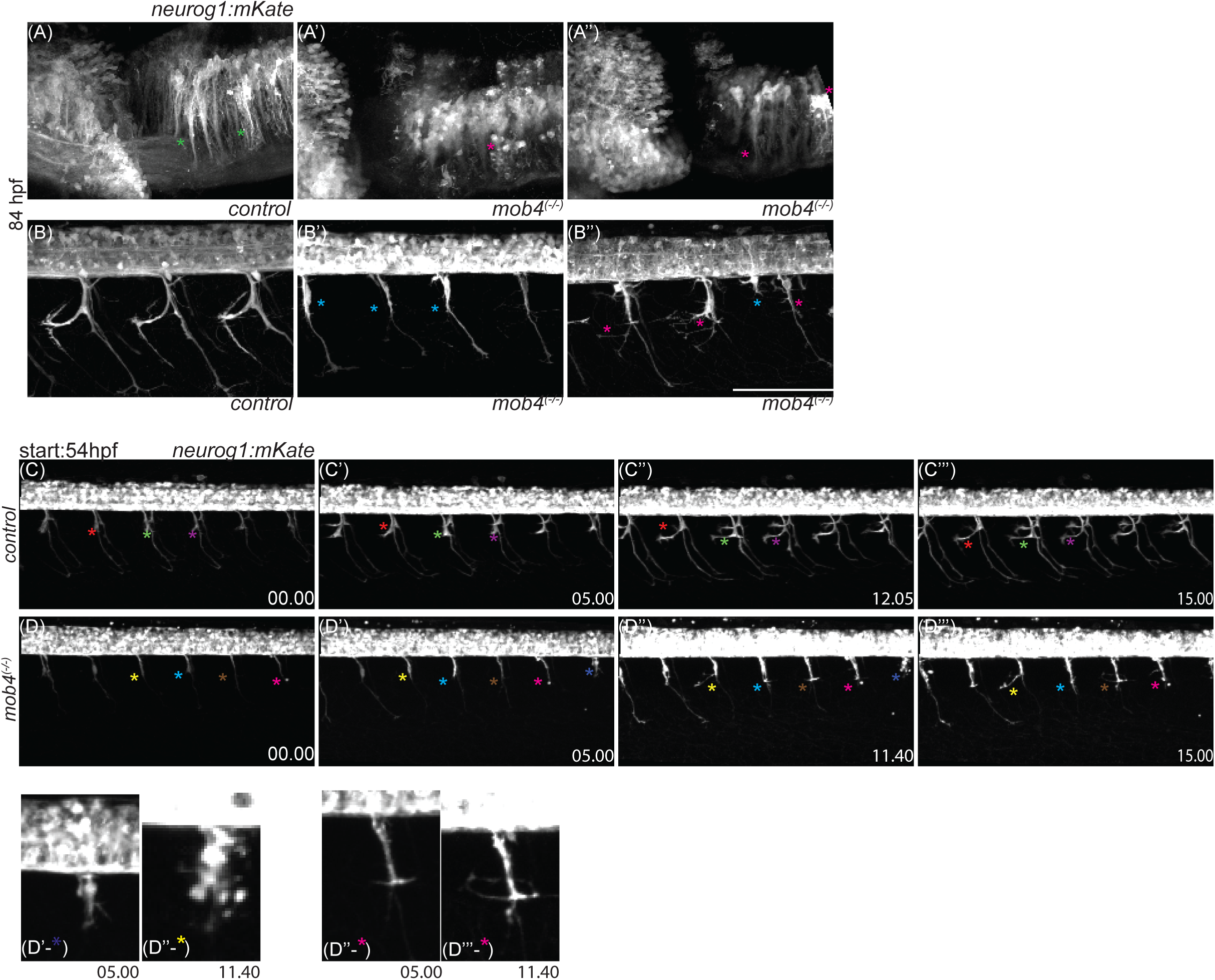
Mob4 lof leads to neuronal defects. (A-A’’) Cranial neurons in *control* (A) and *mob4 mutant* (A’-A’’) 84 hpf embryos. Neuronal bundles are marked with asterisks (WT: green asterisks, aberrant: magenta asterisks). (B-B’’) tail trunk motor neurons in *control* (B) and *mob4 mutant* (B’-B’’m blue asterisks motor neurons with no outgrowth, magenta asterisks MN with hypersprouting) 84 hpf embryos. (A-B’’’) Snapshots of time lapse imaging of development of motor neurons (*ngnl:mKate*) in tail trunk of *control* (C-C’’’) and *mob4 mutant* (D-D’’’) embryos. Time stamps (hh:mm) represent time after start of imaging (00:00). Asterisk colours follow single motor neurons. Selected aberrant (D’-*, D’’-*) neurons are magnified to enhance detail. Scale bar=100µm.

### Continuous requirement of Mob4 function during development and adult physiology

Our analyses indicated a number of cell types affected by Mob4 loss. To further examine cell-autonomous functions of Mob4, we generated a conditional rescue transgenic line (*Tg ubi:loxP-mScarletNLS-loxP-TurboMob4EGFP*) in which Mob4 expression can be activated via temporal- or tissue-specific Cre activity (Fig S2A). A *hsp70l:Cre* transgenic line (Pan et al., 2011) was used to induce rescue transgene expression via dual heat shock at 24 and 30 hpf. While wild-type transgenic embryos appeared normal at 120 hpf (Fig S2C-D), all *mob4* (-/-); *ubi:loxP-mScarletNLS-loxP-TurboMob4EGFP; hsp70l:Cre* embryos without heat shock demonstrated expected phenotypes including cardiac, ocular and brain oedemas, small eyes, defects in body shape and improper fin development (Fig S2E-E’). This indicated minimal leakiness of the rescue transgene. In contrast, *mob4* (-/-); *ubi:loxP-mScarletNLS-loxP-TurboMob4EGFP; hsp70l:Cre* embryos treated with double heat shock showed consistent rescue of phenotypes at 120 hpf in 4/5 embryos, with 1/5 embryos exhibiting a curved body axis (Fig S2 F-F’). These results correspond to previous studies showing ubiquitous expression of Mob4 to rescue loss-of-function defects (Schulte et al., 2010)

Having established that the *mob4* mutant phenotype can be rescued by transgenic Mob4 expression, we next examined the temporal requirements for Mob4 function. We generated a *ubi:loxP-Mob4EGFP-loxP-TagBFP* rescue transgenic line (Fig 6A) which exhibits robust EGFP expression in various tissues (Fig 6-B: brain, B’: endothelial cells, B’’: muscles, B’’’: epithelial cells). In epithelial cells, Mob4EGFP signal is clearly absent from cell membranes (Fig 6B’’’, magenta asterisks), aligning with data from other studies. The Mob4 rescue transgene was crossed into a *mob4* mutant background with a *hsp70l:Cre* transgene for heat-shock-mediated deletion of Mob4 transgenic expression (Fig 6C). In the absence of heat shock, the transgene was sufficient to rescue the *mob4* mutant phenotype in all cases (Fig 6 C’). Following heat shock at 30 hpf, mutant transgenic embryos exhibited typical *mob4* mutant developmental phenotypes at 120 hpf, including pericardial and eye oedemas, uninflated swim bladder, fin defects, and instances of haemorrhaging (Fig 6C’’-C’’’’). Indeed, no viable *hsp70l:Cre* heat-shocked *mob4* mutant transgenic adults were recovered following early heat shock (Fig 6D), demonstrating that Cre-mediated deletion of the rescue construct was robust. Having demonstrated a continuous (post-30 hpf) requirement for Mob4 function in embryonic development, we next asked if Mob4 is required for adult physiology. Adult (6-month-old) *mob4(-I-)*; *ubi:lox-Mob4EGFP-loxTagBFP (hemizygous); hsp70l:cre* fish were subjected to heat shock and subsequently monitored (Fig 6D’). While the majority of *mob4* mutant fish (9/13) died over a period of 5 months post-heat shock, limited mortality was observed in a heterozygous *mob4* mutant transgenic control group (1/6). These results indicate a continuous requirement for Mob4 function in embryonic development and adult physiology.

**Figure 6:**
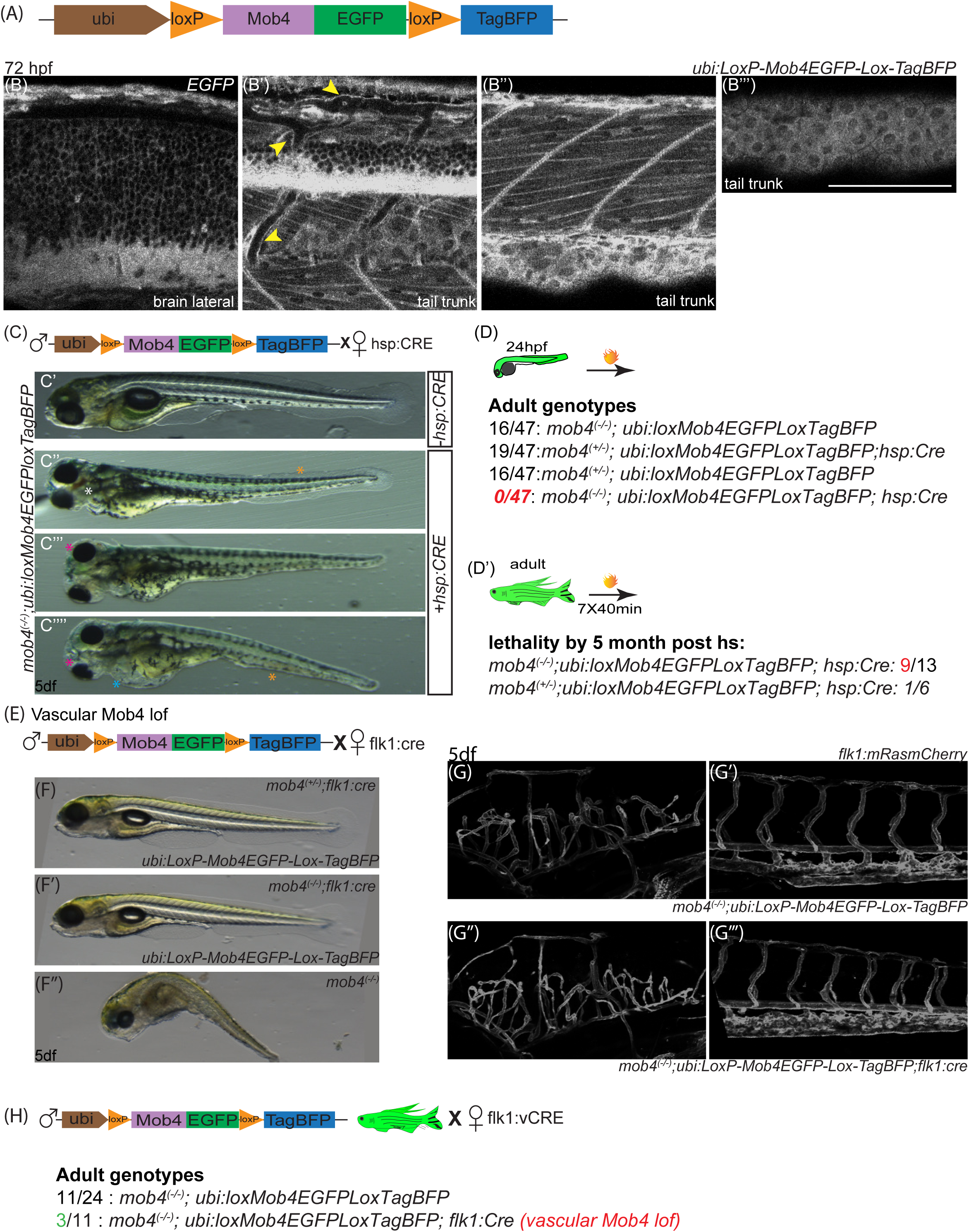
Mob4 has cell specific roles during development. (A) Schematic of rescue transgene. (B-B’’) Localisation of Mob4EGFP in the brain (B), endothelial cells (B’, yellow asterisk), notochord (B’ orange asterisk), epithelial cells (B’’’, magenta asterisk exclusion of Mob4EGFP from cell membrane). (C) Schematic for heat shock induced recombination of *Mob4EGFP* from rescue transgene. (C-C’’’) Heat shocked *mob4^(-I-)^* embryos with rescue transgene, *ubi:loxMob4EGFPloxTAGBFP* (C’) without *hsp70l:Cre* (C’’-C’’’) with *hsp70l:Cre*. (D) Top panel: schematic of experiment to determine long term efficiency of heat shock induced Mob4 loss model. bottom panel: genotypes of resultant adults. (D’) Top panel: schematic of experiment to determine heat shock induced Mob4 loss in adults, bottom panel: score of lethality in *mob4 mutant* vs *control* adults with rescue transgene *ubi:loxMob4EGFPloxTAGBFP*. All adult fish are positive for *hsp70l:Cre*. (E) Schematic for Vascular Mob4 loss model. *kdrl:Cre* induced recombination of *Mob4EGFP* from rescue transgene *ubi:loxMob4EGFPloxTAGBFP* in endothelial cells. (F-F’’) early development in endothelial Mob4 loss model. Representative brightfield images of whole embryos at 5 dpf. (F) *mob4^(+I-)^*;*ubi:loxMob4EGFPloxTAGBFP;kdrl:Cre*, *control* (F’) *mob4(-I-)*;*ubi:loxMob4EGFPloxTAGBFP*;*kdrl:Cre*, vascular Mob4 loss (F’’) *mob4 mutant* embryo, global mob4 loss. (G-G’’’) vascular development in vascular Mob4 loss model with *kdrl:hRasmCherry* labelled vasculature at 5dpf. (G, G’) brain and tail trunk vasculature in *mob4(-I-)*;*ubi:loxMob4EGFPloxTAGBFP* (control, rescued embryos). (G’’-G’’’) brain and tail trunk vasculature in *mob4(-I-)*;*ubi:loxMob4EGFPloxTAGBFP;kdrl:Cre* (vascular Mob4 loss). (H) Left panel: Schematic to determine effect of vascular Mob4 loss in adults. Right panel: genotyping of adults. Scale bar=100μm.

We next examined the function of Mob4 in endothelial cells via the use of our conditional rescue transgenic and a *kdrl:cre* EC-specific transgenic (Fig 6E). In *mob4(-I-); ubi:loxP-Mob4EGFP-loxP-TAGBFP; kdrl:hRasmCherry* embryos without the *kdrl:cre* transgene (Fig S2 A-A’’), no signal was observed for Tag-BFP (Fig S2A). hRasmCherry labels all endothelial cells (Fig S3A’). Mob4GFP was present in endothelial cells (yellow arrows S2A’’). In *mob4(-I-); ubi:loxP-Mob4EGFP-loxP-TAGBFP; kdrl:hRasmCherry; kdrl:cre*, Cre was sufficient to activate robust TagBFP in the vasculature (Fig S3B) with simultaneous reduction of GFP in vascular cells (Fig S2B’-B’’ orange asterisks). Interestingly, *mob4(-I-); ubi:loxP-Mob4EGFP-loxP-TAGBFP; kdrl:Cre; kdrl:hRasmCherry* embryos appeared superficially normal at 120 hpf, with an absence of observable cranial hemorrhaging or cardiac oedemas (Fig 6F-F’’) and normal vascular patterning at 120 hpf (Fig 6G-G’’’). Transgenic (GFP-positive) embryos from these experiments were raised to adulthood and subsequently genotyped (Fig 6H). Of the *mob4* mutants recovered (11/24 fish), three were additionally carriers of the *kdrl:Cre* transgene, suggesting that Mob4 loss in endothelial cells alone may be insufficient for severe or lethal phenotypes. Our results indicate that although global Mob4 loss leads to a spectrum of defects in cardiovascular and neuronal development, Mob4 has pleiotropic tissue-specific roles during development and adult life.

## Discussion

In this study, we have used the loss of Mob4 function in zebrafish as a proxy for examining the role of the STRIPAK complex in vertebrate development, in particular vascular development. Our proximal interactome of Mob4 in early zebrafish embryos confirms *in vitro* work (Goudreault et al., 2009; Jeong et al., 2021; Kean et al., 2011) defining Mob4-STRIPAK associations. As both Mob4 and Striatin3 loss led to similar defects in muscle and neuronal development in zebrafish (Berger et al., 2022), these data argue that Mob4 is an integral component of STRIPAK in zebrafish. We show that Mob4 loss leads to the eventual cessation of blood flow in zebrafish, with defects in cardiac and vascular development. Notably, *mob4* mutants demonstrate both intracranial haemorrhages at sites where angiogenesis was at first apparently normal, and venous hypersprouting in the trunk vasculature. Taken together, these results argue for Mob4/STRIPAK function regulating the stability of developing vascular networks (at least in the case of the tail trunk vasculature) in an endothelial cell-specific manner.

The signalling pathways that Mob4/STRIPAK act through and that are dysregulated in *mob4* mutants remain to be determined. Components of the Wnt signalling pathway, including Axin complex tumour suppressor adenomatous polyposis coli gene product APC (Apc), and dishevelled segment polarity protein 3a (Dvl3a), are enriched in our Mob4 TurboID dataset. Other enriched proteins relate to the neuronal phenotypes observed in *mob4* mutants, including Filamin-interacting protein Filip1 and the atypical phosphatase Dusp27. Filip1 regulates dendritic spine development through modulation of non-muscle myosin 2b (Yagi et al., 2014), while Dusp27 is required to establish proper Z-disc structure (Chen et al., 2023). As Mob4 is also known to regulate cytoskeleton components in various systems, our data raises the possibility that it acts through these and other regulators of cytoskeletal development.

Although Mob4 has been shown to be enriched in various neuronal and glial tissues, current data suggest that Mob4 has a broader role in organogenesis (Berger et al., 2022; Schulte et al., 2010). Our mutant, rescue, and conditional loss-of-function models provide tools to understand the tissue-specific functions of Mob4. The rescue of trunk venous vasculature defects by *Lyve1:Mob4* strongly suggests an endothelial cell-specific function for Mob4. In contrast, *kdrl:Cre* (pan-EC)-mediated loss of Mob4 resulted in no discernible phenotypes. Unfortunately, the nature of the transgenic elements used for these experiments precluded being able to determine if Mob4 was completely lost in ECs, although we did observe (via expression of TAGBFP) robust recombination of the rescue transgene in ECs (Fig S3B). Given that incomplete Cre-mediated excision is observed for many zebrafish transgenic lines (Juan et al., 2024; Lalonde et al., 2022), and the possibility that multiple rescue transgene copies were present in transgenic lines used for these studies, it is possible that Mob4 was only lost in a mosaic manner in these experiments. It should be noted that *mob4* mutants, due to cardiac defects, lose blood flow over time, which in turn leads to well-characterized cardio-vascular defects (Kugler et al., 2021; Sehnert et al., 2002). The major vascular phenotypes observed in this study – cerebral haemorrhages and venous hypersprouting – have not been reported with reduced or absent blood flow in zebrafish. Interestingly, *mob4* mutant embryos exhibited haemorrhaging only in the head and not in the tail trunk vasculature, suggesting a specific function of Mob4 in maintaining the blood-brain barrier. Given the prominent neuronal defects observed in *mob4* mutants, and the role of secreted neuronal factors in cranial vasculature integrity (Hartmann et al., 2023), it is possible that Mob4 functions in multiple (EC, neuronal and perhaps other) cell types. More robust Cre deletor lines will be required to examine this, as the *hsp70l:Cre* transgenic appeared to be sufficient to greatly reduce Mob4 rescue expression.

Overall, we have shown that Mob4 functions as a component of the STRIPAK complex in zebrafish and has tissue-specific roles in the regulation of early cardiovascular and neuronal development. The latter, essential roles of Mob4 in adult physiology remain to be uncovered. However, the conditional rescue and loss of function tools we have developed, coupled with genetic and proteomic approaches, will, in the future, elucidate critical aspects of Mob4 and STRIPAK function.

## Materials and Methods

### Zebrafish and line maintenance

Adult AB/TL mixed strain zebrafish (Danio rerio) were maintained as per Canadian Council on Animal Care (CCAC) and The Hospital for Sick Children Animal Services (LAS) guidelines. The following previously described lines were used in our experiments: *Tg[kdrl:EGFP]* (*s843*); *Tg[gata1:dsred]* (*sd2*), *Tg[pdgfrb:Citrine]* (*s1010*), *Tg[tp1-MmHbb:EGFP]* (*um14*), *Tg[kdrl:hRas-mCherry]* (*s896*), *Tg[hsp70l:Cre]* (*zdf13*), *Tg[ngnl:Kate]* (*sk100*), *Tg[kdrl:Cre]* (*s898*).

### Generation of Crispr/Cas9 mutant line

The *mob4* (hsc205) mutant line was generated via use of CRISPR/Cas9 as described previously (Jao et al., 2013). Cas9 was generated as previously described (Gagnon et al., 2014). gRNA-Cas9 complex was injected High resolution melt (HRM) (Parant et al., 2009) was used to test the efficacy of mutants with the following primers: **F**:CGACGAAAGATCGTTATATGGTT, **R**:ATGAAGGACCCTTGTTGACATT Subsequent genotyping for the *mob4* allele harbouring an 11bp deletion resolvable on an agarose gel was carried out with PCR using the same primers.

### Plasmids and generation of transgenic lines

Gateway cloning was caried out with Gateway™ LR Clonase™ II Enzyme mix, and Gateway™ BP Clonase™ II Enzyme mix from Thermos Fisher and various plasmids from the Tol2kit (Kwan et al., 2007).

gRNA Plasmid: oligonucleotides TAGGAGATTATGGAGCCTCCAG and AAACCTGGAGGCTCCATAATCT were annealed and cloned into pT7-gRNA with BslI. gRNA was generated with BamHI/T7 MEGAshortscipt T7 (Invitrogen).

***pCS2+Mob4***: The entire *mob4* CDS was amplified from 48 hpf *WT* cDNA with **XhoI_mob4_F**: tatatCTCGAGatggtcatggc and **Mob4_XbaI_R**: tatataTCTAGAtcaggcatcgct and cloned into pCS2+ with XhoI and XbaI.

***pCS2+turboMob4EGFP****: mob4* was amplified *pCS2+Mob4* with **XhoI_mob4_F** and **Mob4EGFP_OL_R**: TGCTCACCATGGCATCGCTT and *EGFP* was amplified with **Mob4EGFP_OL_F**: AAGCGATGCCATGGTGAGCA and **XbaI_EGFP_R**:

TATATATCTAGATTACTTGTACAGCTCGT. Fusion PCR was carried out on the PCR products with **XhoI_mob4_F** and **XbaI_EGFP_R**, to generate *mob4EGFP*, digested with XhoI and XbaI and cloned into pCS2+ to generate *pCS2+mob4EGFP*. *Turbo* was amplified with **XhoI_turbo_F**: tataCTCGAGGCAAACATGGCGAAAGATAAC and **turbo_mob4_OL_R**: atgaccatAGATCCTCCCCC from PCS2+turbo (Rosenthal et al., 2021) *mob4EGFP* was amplified with **turbo_mob4_OL_F**: GGGGGAGGATCTATGGTCAT and **XbaI_Mob4_R**: tatataTCTAGAtcaggcatcgct from *pCS2+mob4EGFP*. *Turbomob4EGFP* was amplified with fusion PCR with **XhoI_turbo_F** and **XbaI_Mob4_R** and cloned into pCS2+ with XhoI and XbaI. Capped *Turbomob4EGFP* mRNA was generated with SP6 mMessage mMachine kit (ThermoFisher).

***pME:TurboMob4EGFP***: *TurboMob4EGFP* was amplified from *pCS2+Turbomob4EGFP* with **attb1_turbo_kozak**_**F**:ggggacaagtttgtacaaaaaagcaggctaagcaaacatggcgAAAGATAAC and **attb2-EGFP-R**:ggggaccactttgtacaagaaagctgggtaTTACTTGTACAGCTCGTC and cloned into pME with BP Clonase.

***pDest[fli1a:mApple,cmlc2EGFPJ***: Gateway cloning with plasmids from the. *p5E-fli1a* (Lawson Lab), *pME:mApple* (Tol2kit) and *p3E:polyA* (Tol2kit) were gateway cloned with LR Clonase (Gateway™ LR Clonase™ II Enzyme mix) into *p0estCG2* (Tol2kit) to generate pDest*[fli1a:mApple,cmlc2EGFP]*.

***pDest[lyve1:Mob4EGFPJ***: p5E:Lyve1 (Tol2Kit), pME:Mob4EGFP, and p3E:polyA (Tol2kit) were cloned into destination vector pDEst-CG2 (Tol2kit)

***pDest[ubi:lox mSacrletNLS lox TurboMob4EGFPJ***: the simpleswitch parent plasmid *p0est[ubi:lox-mSacrletNLS-loxWasabi]* was from a previously published study (Burgess et al., 2020). *TurboMob4EGFP* was amplified from *pCS2+TurboMob4EGFP* with **Sbf1_Turbo_F**: tataCCTGCAGGgcaaacatggcgAAAGATAACand and **AscI_EGFP-STOP_rev**: tataggcgcgccTTACTTGTAC, digested with SbfI and AscI, and used to replace wasabi in the simple switch plasmid to generate *p0est[ubi:lox-mSacrletNLS-loxTurboMob4EGFP]*.

***pDest[ubi: lox Mob4EGFP lox TagBFPJ***. *Tag-BFP* was amplified with **Sbf1_Tag-BFP_F**: tataCCTGCAGGgcaaacatggtgagcgagc and **AscI_BFP_Rev**: tataGGCGCGCCttaattaagctt and digested with SbfI and AscI and used to replace Wasabi in the Simpleswitch parent plasmid to generate *p0est[ubi:lox-mSacrletNLS-loxTagBFP]*. Next M*ob4EGFP* was amplified with **AgeI_Mob4_F:** tataaccggtgcaaacatggtcatggcggagg and **AgeI_EGFP_R**: tataaccggtTTACTTGTACAGC and used to replace mScarletNLS between the lox sites in *p0est[ubi:lox-mSacrletNLS-loxTagBFP]* to generate *p0est[ubi:lox-Mob4EGFP-loxTagBFP]*.

Transgenic lines were generated as described (Suster et al., 2011). Briefly 20-50 pg *p0est* plasmid DNA and 100pg *tol2* mRNA was co-injected in single cell stage embryos. Embryos with robust mosaic expression of selection markers were grown up and screened for F1 germline transmission to establish stable lines. The following hsc numbers are associated with the lines generated in this work: *hsc201*:*Tg[fli1a:mApple,cmlc2:EGFP]*, *hsc202*: *Tg[lyve1:Mob4:EGFP]*, *hsc203*: *Tg[ubi:lox mScarletNLS lox TurboMob4EGFP]*, *hsc204*: *Tg[ubi: lox Mob4EGFP lox TagBFP]*. Plasmids for the generation of these lines or the lines themselves are available upon request to ICS. The *Tg[mrc1a:EGFP] line* was generated from a plasmid generously provided by the lab of Dr. Brant Weinstein and has been previously described (Jung et al., 2017).

### Microscopy and image analysis

Whole embryo images were acquired using Zeiss AXIO Zoom V16. Confocal images were acquired with Nikon A1R laser-scanning confocal microscope using either 20X or 40X water immersion objectives. Images were analysed and processed with FijI (Schindelin et al., 2012). Embryos were mounted in 0.8% low melt agarose in egg water in clear bottomed round petri dishes, covered with embryo media, and placed on a heated plate holder at the microscope for all live imaging. These embryos were carefully removed from agarose at the end of imaging and genotyped as required.

### Proximity Dependant Biotinylation (BioID)

BioID experiments followed the protocols as described in Rosenthal et al. 2021. To express the TurboID fusion proteins 200pg of in vitro transcribed *TurboMob4EGFP* mRNA was injected into single cell stage embryos. The embryos were incubated in egg water supplemented with 800 µM biotin from 12 hpf. At 48 hpf labelling was terminated by transfer to ice cold egg water. The embryos were then deyolked and frozen at -80°C. For protein extraction and pulldown, the embryos lysis and streptavidin affinity purification were performed as described. Proteins were prepared for mass spectrometry analysis as described. Mass spectrometry acquisitions were performed using a ThemoFisher Fusion Lumos mass spectrometer as described. Mass spectrometry data analysis was performed as described. Briefly, ProteoWizard (Kessner et al., 2008) was used to convert the .RAW files. The files were searched using Mascot (Perkins et al., 1999) and Comet (Eng et al., 2013) against zebrafish sequences from the RefSeq database. The results from each search engine were analysed with the Trans-Proteomic Pipeline (Deutsch et al., 2010) via the iProphet pipeline (Shteynberg et al., 2011). Proteins with an iProphet probability ≥95% and two unique peptides were used for analysis. For interaction scoring the SAINTexpress (Teo et al., 2014) tool was used to score the probability of identified proteins being enriched above background. Briefly, SAINTexpress uses the number of spectral counts for each prey identified in the negative control runs compared against the average from the two replicates of the experimental runs to calculate a Bayesian False Discovery Rate (BFDR); preys with a BFDR ≤5% were considered high confidence. All data analysis was performed within the ProHits laboratory information management system (LIMS) platform (Liu et al., 2010). For data visualization, dot plots were generated using ProHits-viz (Knight et al., 2017) . To perform Gene ontology gene set enrichment analysis human orthologs of the identified high confidence proteins were retrieved using the DIOPT online tool (Hu et al., 2011). This conversion was performed to take advantage of the much greater number of annotations available for human genes than for the zebrafish. Gene ontology gene set enrichment analysis was performed using the g:Profiler g:GOSt tool (Reimand et al., 2011) with default settings.

## Acknowledgements

We wish to thank members of the Gingras and Scott labs for helpful discussions during the execution of these experiments and writing of this manuscript. We would also like to thank Dr. Brent Derry for valuable input during manuscript preparation and Dr. Brant Weinstein for providing us with his *Tol2(mrc1a:EGFP)* construct. All zebrafish experiments were performed in the zebrafish facility at The Hospital for Sick Children following the guidelines of the institutional Animal Care Committee and the Canadian Council on Animal Care. We thank Matthew Carpenter, Tyler Cunningham, Solange Prokop, Alejandro Salazar and their associated work study students for expert care of animals used in this study. Proteomics work was carried out at the Network Biology Collaborative Centre at the Lunenfeld-Tanenbaum Research Institute by Brett Larsen and Cassandra Wong. SMR was supported by Hospital for Sick Children Restracomp and Ontario Graduate Scholarship studentships.

This work was supported by funding from the Natural Sciences and Engineering Council of Canada (NSERC, RGPIN-2017-06502 to ICS) and the Canadian Institutes of Health Research (CIHR, PJT 153000 to ICS).

---

**Figure S1:**
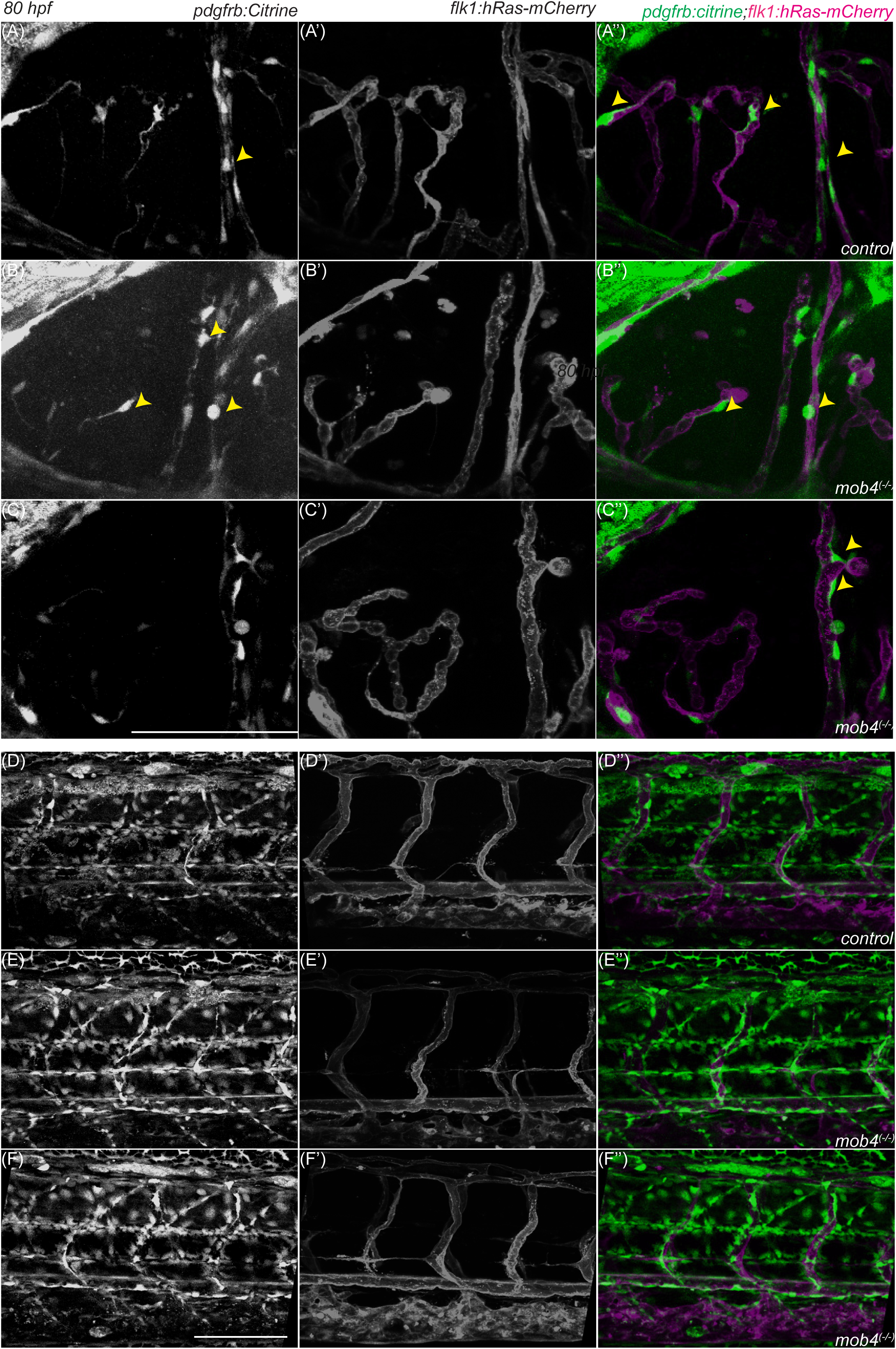
Pericyte localisation in vasculature of *mob4 mutant* embryos. *pdgfrb:Citrine* labels pericytes with Citrine and *kdrl:hRasmCherry* labels all ECs. (A-C’’) Projections of confocal z-stacks of head (A-C’’) and tail trunk (D-F’’) of 80 hpf embryos imaged laterally. (A-A’’, D-D’’) *control* (B-C’’, E-F’’) two representative *mob4 mutant* embryos each for head and tail trunk vasculature. Pericytes associated with ECs are indicated with yellow arrowheads. Scale bar=100μm

**Figure S2:**
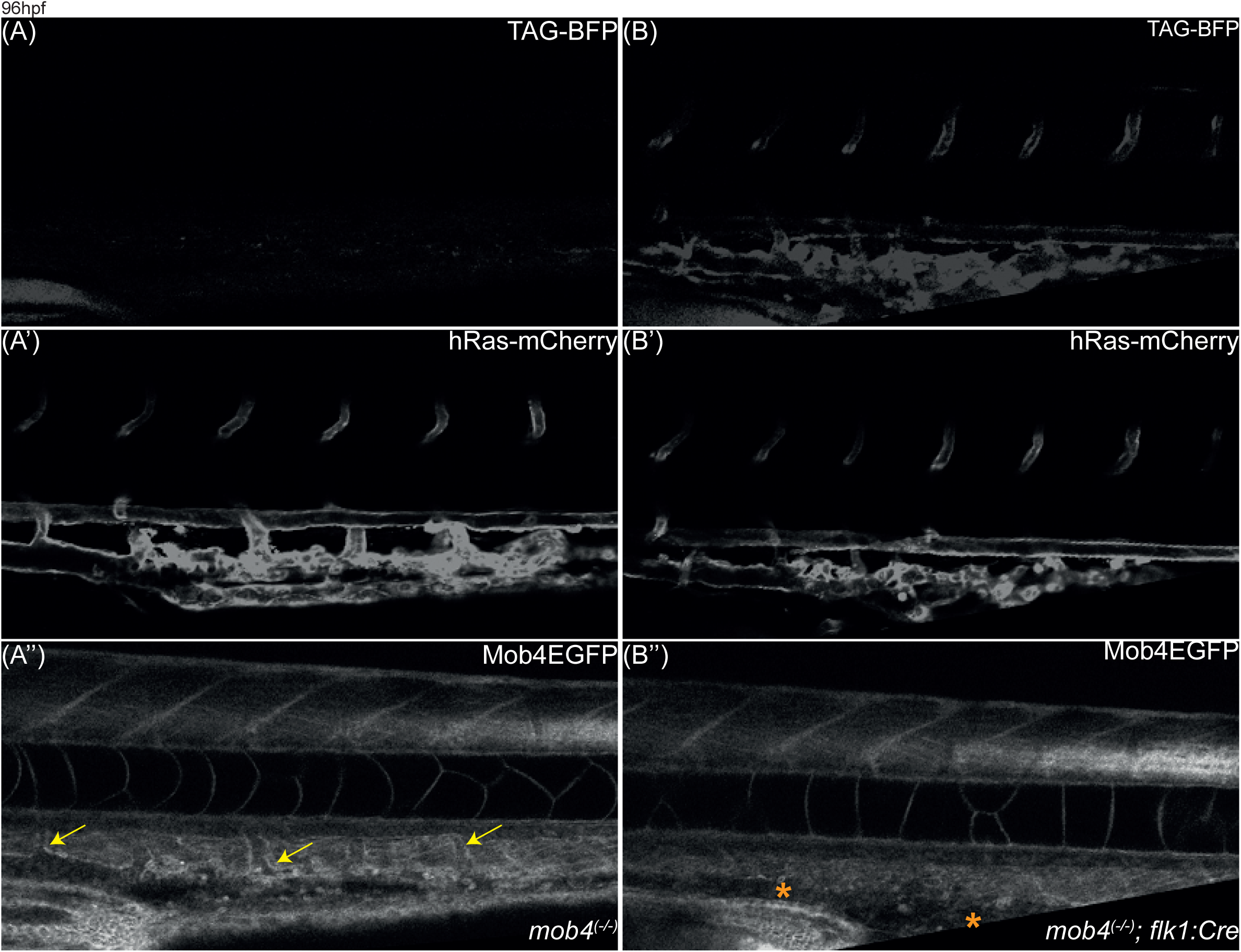
Action of Switch Transgene in the vasculature. (A-B’’) Confocal images of vasculature in the tail trunk. (A-A’’) *mob4(-I-);ubi:loxMob4EGFPlox TagBFP* (B-B’’) *mob4(-I-);ubi:loxMob4EGFPlox TagBFP; kdrl:Cre.* Scale bar=100µm

**Figure S3:**
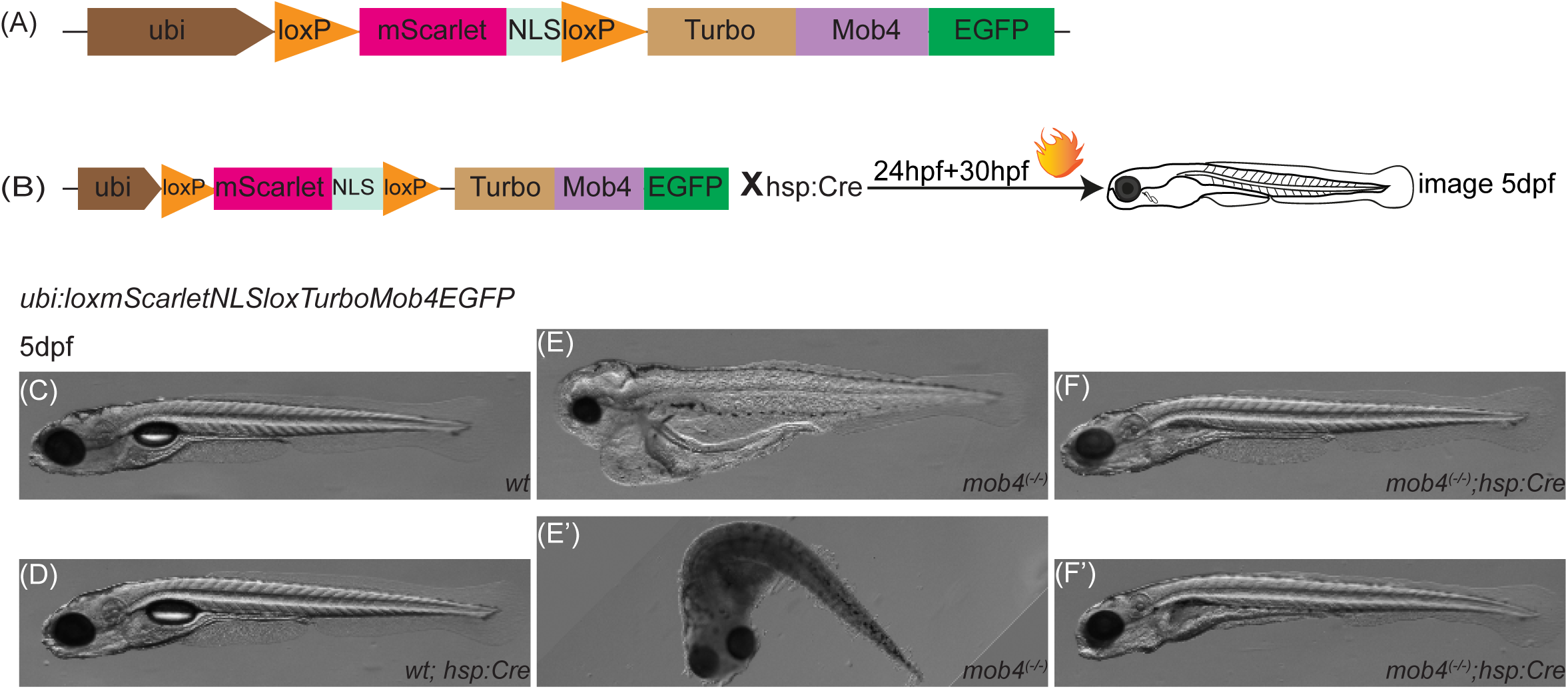
(A) Schematic for rescue transgene (B) Schematic of heat shock dependent rescue experiment. Heat shock carried out at 20 hpf and 30 hpf. Whole embryos imaged at 5dpf. All embryos C-F’ are positive for *ubi:loxP-mScarletNLS-LoxP-TurboMob4EGFP* (C) only ubi:loxP-mScarletNLS-loxP-TurboMob4EGFP (D) ubi:loxP-mScarletNLS-loxP-*TurboMob4EGFP;hsp70l:Cre* (E-E’) *mob4(-I-);ubi:loxP-mScarletNLS-loxP-TurboMob4EGFP* (F-F’) *mob4(-I-);ubi:loxP-mScarletNLS-loxP-TurboMob4EGFP;hsp70l:Cre* rescued embryos

**Supplementary Movie1**: Time lapse imaging of lateral head vasculature starting at 54 hpf in *kdrl:EGFP*;*gata1:dsRed* double transgenic embryos. EGFP labels all ECs and dsREd (A) *control* (B-D) *mob4 mutant* embryos. GFF. Asterisks show instances of haemorrhage in B and D and vascular anomalies in (C). Scale bar=100μm.

**Supplementary Movie 2**: Time lapse imaging of lateral tail trunk vasculature starting around 54 hpf in *kdrl:EGFP*;*gata1:dsRed* double transgenic embryos (A) *control* (B-D) *mob4 mutant* embryos. Asterisks show aberrantly sprouting ECs. Scale bar=100μm.

**Supplementary Movie 3**: Time lapse imaging of lateral tail trunk vasculature starting around 56 hpf in *tp1:EGFP*; *kdrl:hRAsmCherry* double transgenic embryos. (A-A’’) *control* (B-B’’, C-C’’) *mob4 mutant* embryos. Asterisks label hypersprouting ECs positive for mCherry and negative for EGFP in *mob4 mutant* embryos

**Supplementary Movie 4**: Time lapse imaging of motor neurons in the lateral tail trunk starting around 54 hpf in *ngn1:mKate* transgenic embryos. (A) *control* (B) *mob4 mutant*. Asterisks marks irregularities in neuron outgrowth. Scale bar=100μm.

## Notes

### Competing Interest Statement

The authors have declared no competing interest.

### Summary of Updates

Supplementary Movies and Tables have been included.

